## Supplementary Material for "Machine Learning based histology phenotyping to investigate epidemiologic and genetic basis of adipocyte morphology and cardiometabolic traits"

### Supplementary Information

#### Supplementary Figures

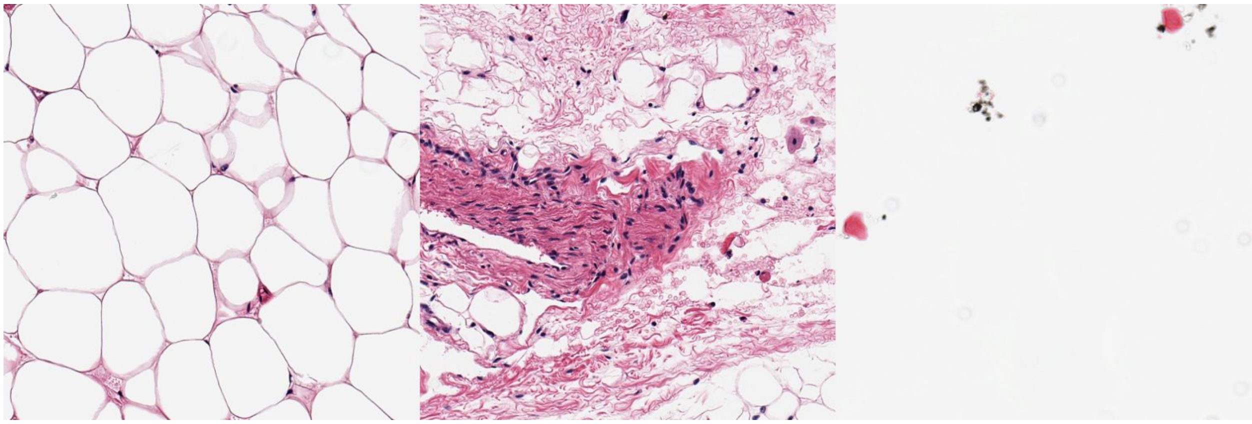

**Supplementary Figure 1 | Example histology imaging tiles.** From left to right. Examples of Adipocyte, non-adipocyte and empty tiles used for initial CNN classifier to propose Regions of Interest (ROIs) of Whole Slide Image (WSI) histology slides.

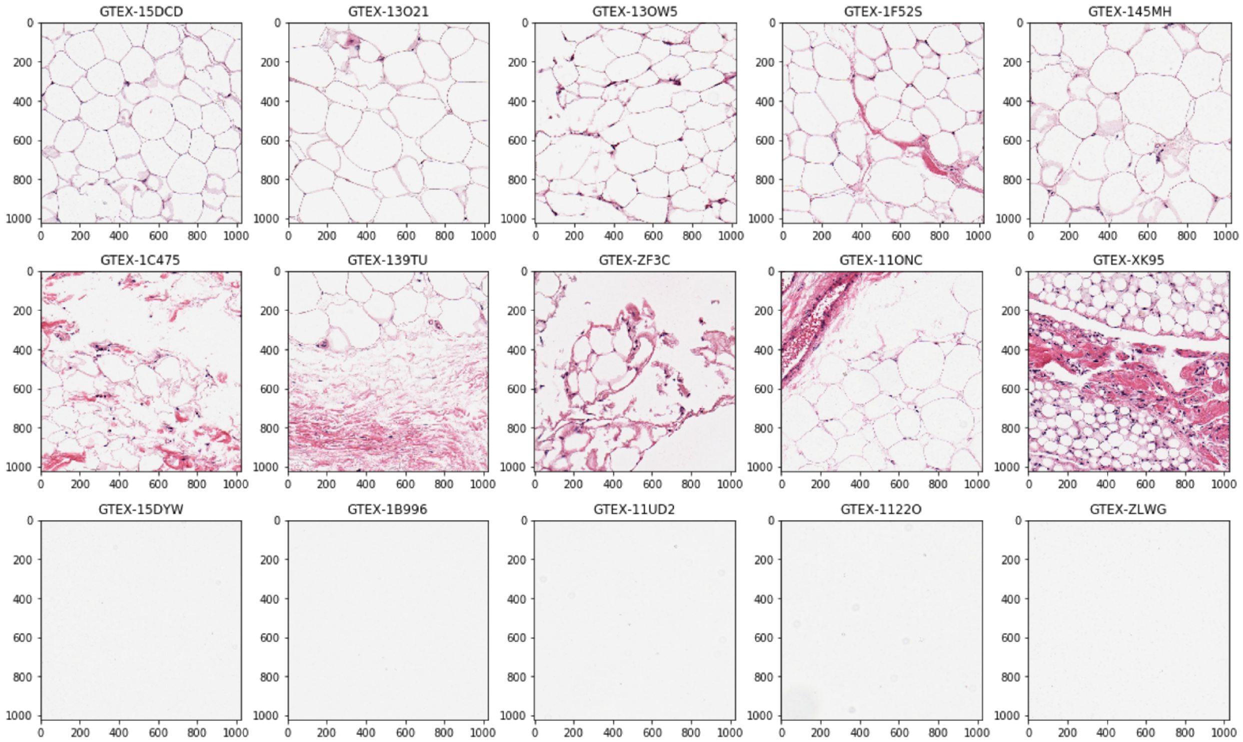


**Supplementary Figure 2 | Examples of classified tiles with Posterior probability $\geq$ 0.9.** Top row: 'Adipocyte tiles', middle: 'not adipocyte', bottom: 'empty'.

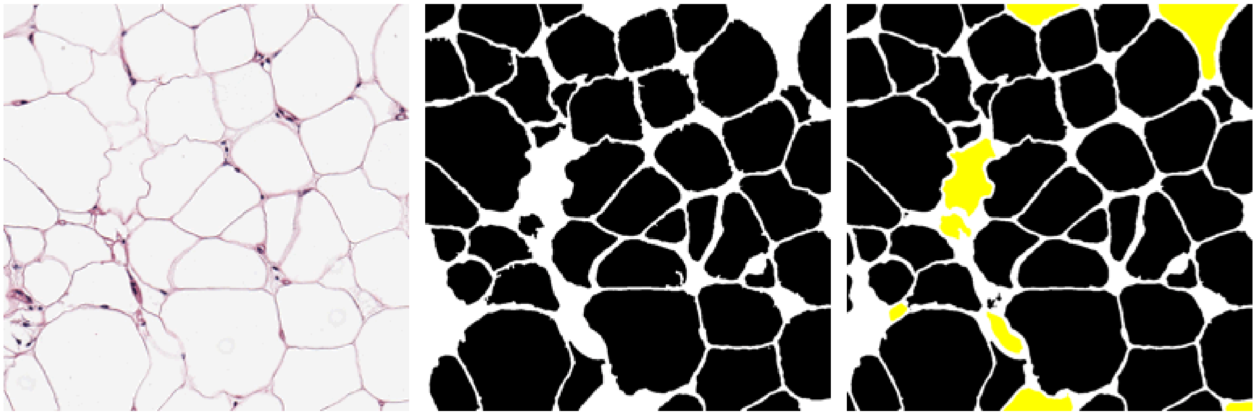

**Supplementary Figure 3 | Adipocyte U-net performance.** From left to right: an out-of-sample adipocyte input tile. Middle: Adiposoft input mask (settings: cell diameter range 20-300$\mu m$). Right: Adipocyte U-net predicted mask after Otsu thresholding. The U-net prediction includes cells missed by Adiposoft - *highlighted in yellow*. Adiposoft counts 45 cells, U-net counts 53 - manual count is 56.

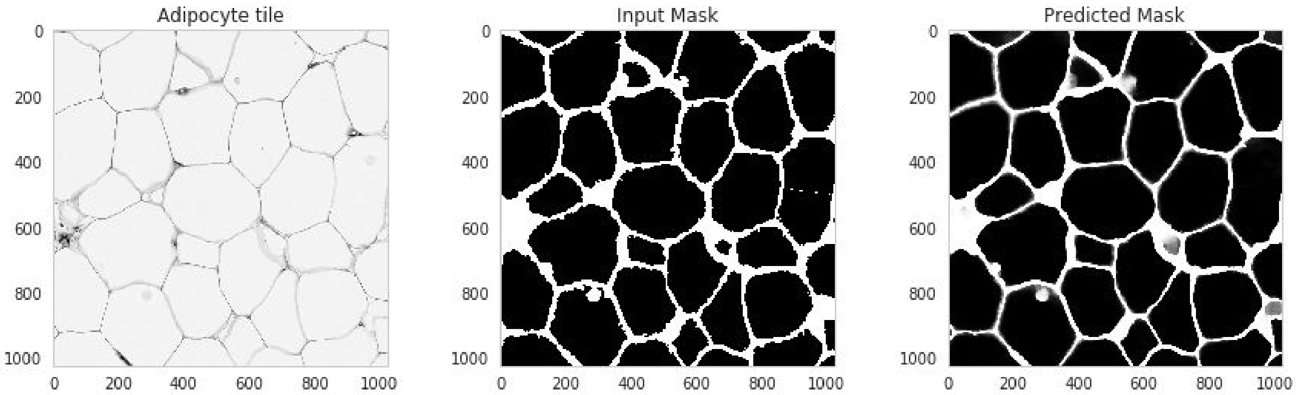


**Supplementary Figure 4 | Example test tile prediction.** From left to right: Input tile, Input mask, Predicted mask and Pixel wise classification results. Predictions highly accurate and recover some cells missed in input mask.

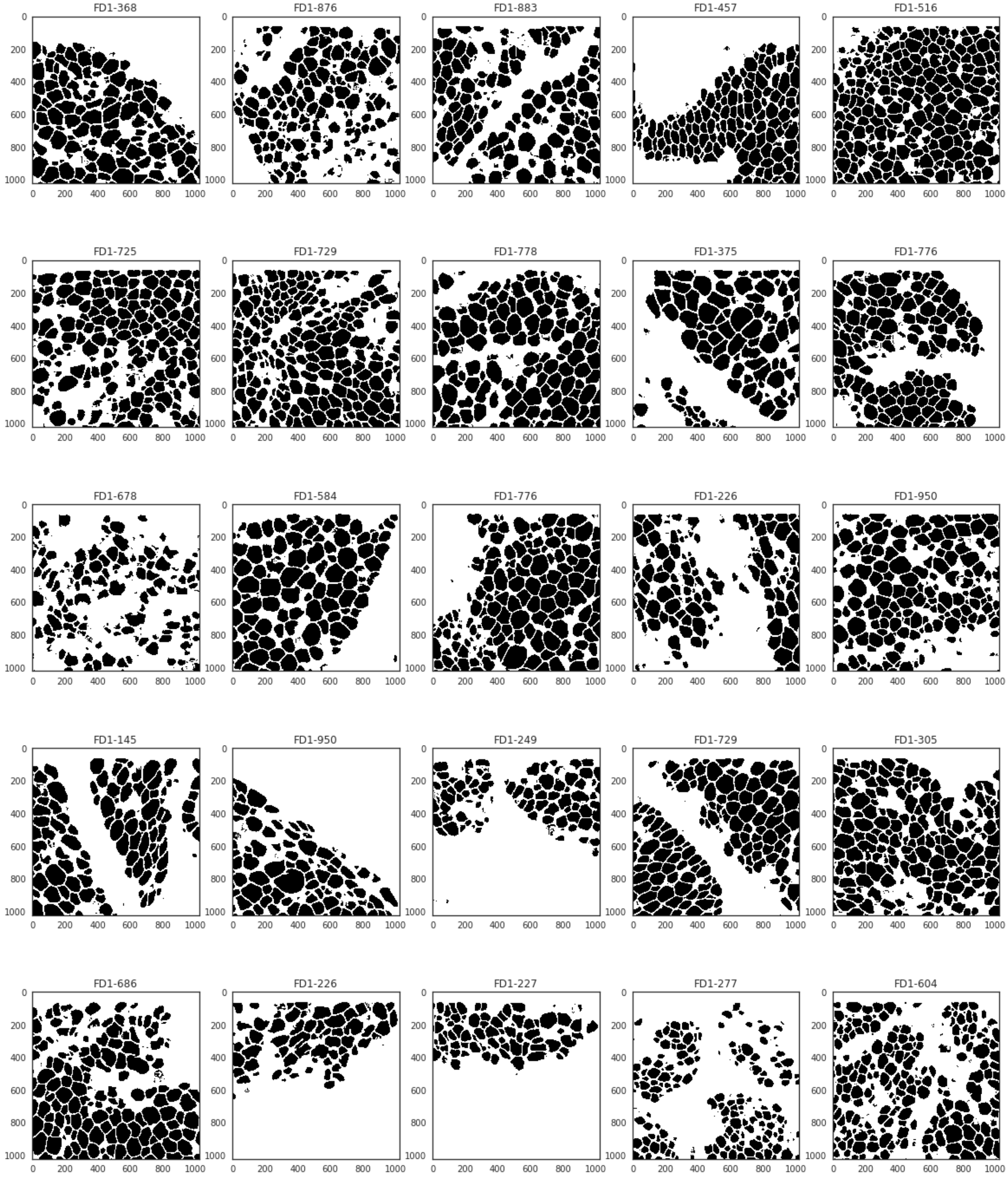


**Supplementary Figure 5 |** 25 random Adipocyte U-net segmentations from the fatDIVA cohort. All segmentations from all cohorts are publicly available through Github.

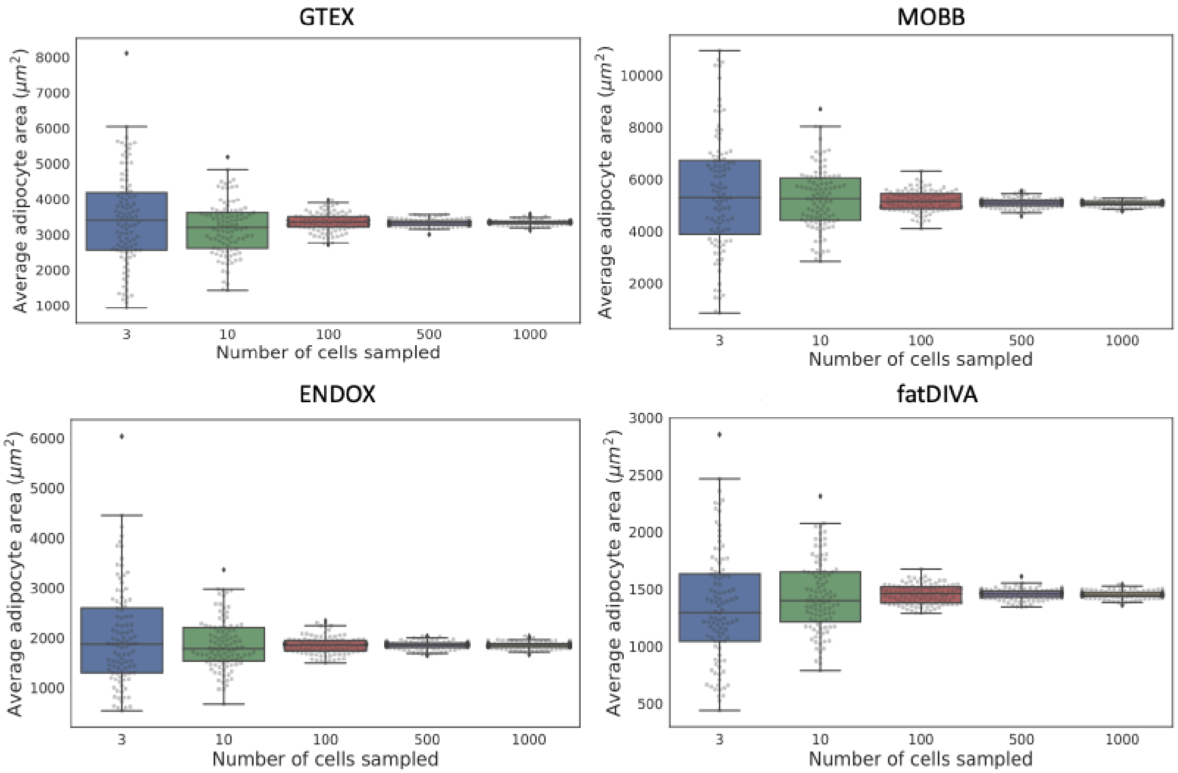


**Supplementary Figure 6 | Adipocyte measures from an individual in each cohort that had > 1000 cells measured.** We randomly sampled 3, 10, 100, 500 and 1,000 cells one hundred times to assess how variability in adipocyte area mean was affected. We determined 500 cells per sample was a suitable number to minimise variance due to subsampling cells.

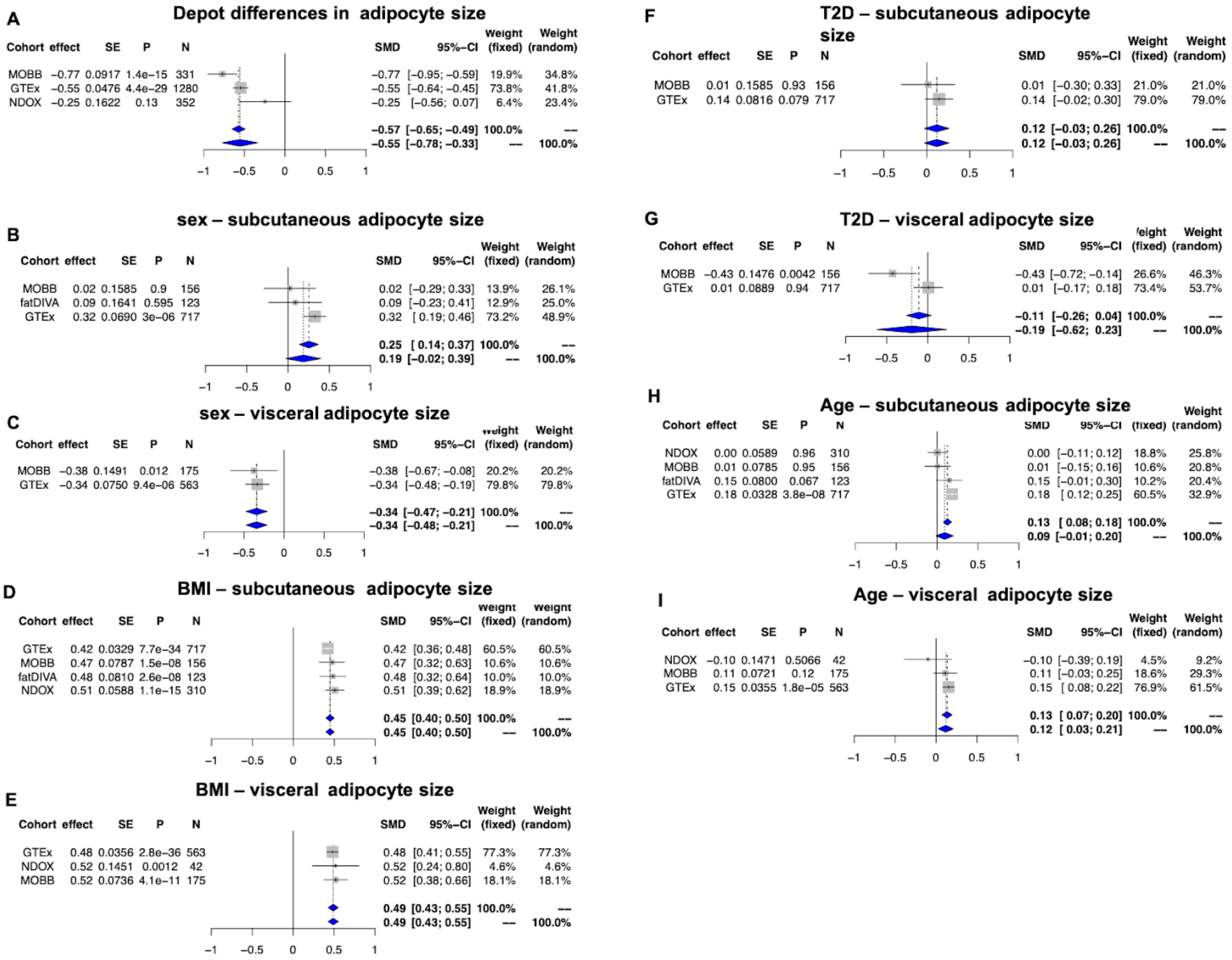

**Supplementary Figure 7 | Phenotype- adipocyte size random effects meta-analysis forest plots**. **A**) Depot specific differences in adipocyte size across cohorts. **B**) Sexual dimorphism in subcutaneous depot adipocyte size across cohorts. **C**) Sexual dimorphism in visceral depot adipocyte size across cohorts. **D**) Association with subcutaneous depot adipocyte size and BMI across cohorts. **E**) Association with visceral depot adipocyte size and BMI across cohorts. **F)** T2D association with subcutaneous depot adipocyte size. **G)** T2D association with visceral depot adipocyte size. **H**) Chronological age association with subcutaneous depot adipocyte size across cohorts. **I**) Chronological age association with visceral depot adipocyte size across cohorts.

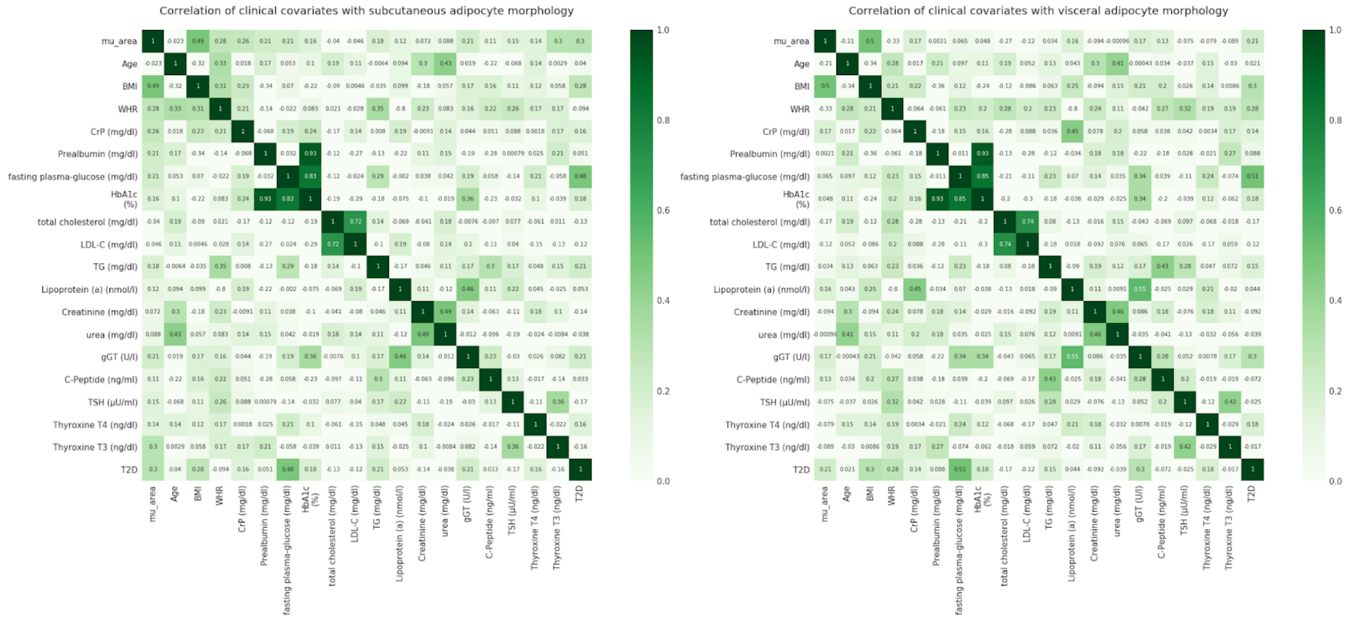


**Supplementary Figure 8 | Correlation between clinical phenotypes in the MOBB cohort and estimated adipocyte surface area measures.** The MOBB cohort contained a large number of covariates related to obesity and cardiometabolic traits. We therefore estimated correlation between adipocyte size and these covariates. In both depots, adipocyte size correlates most strongly with BMI.

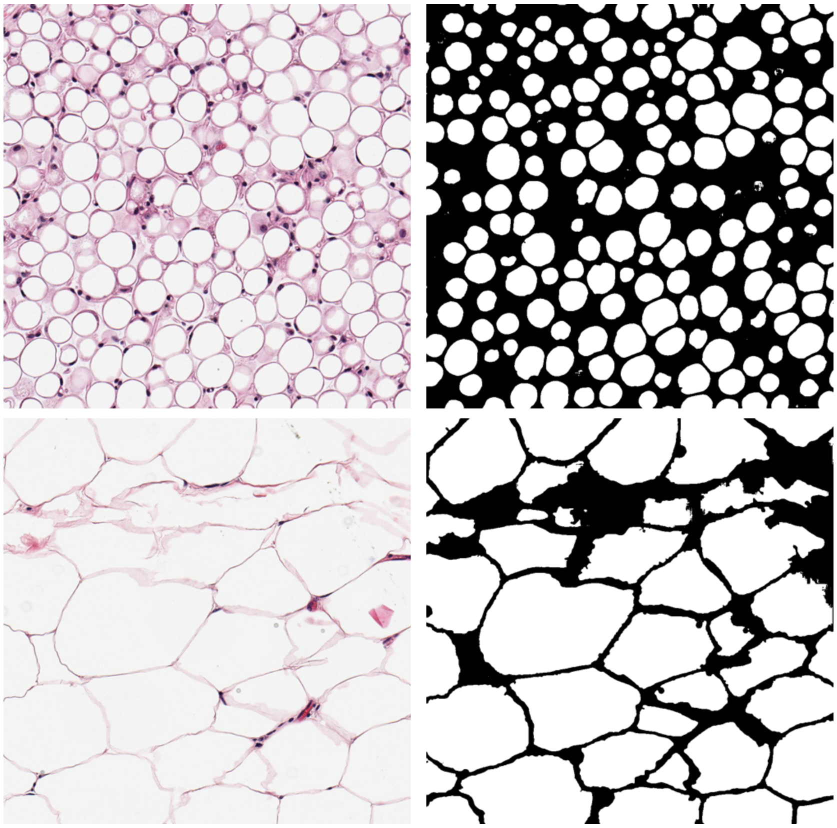

**Supplementary Figure 9** | Example of low adipocyte area variance (top) and High variance (bottom) samples (left) and their corresponding predicted masks (right) - Both Omentum samples.

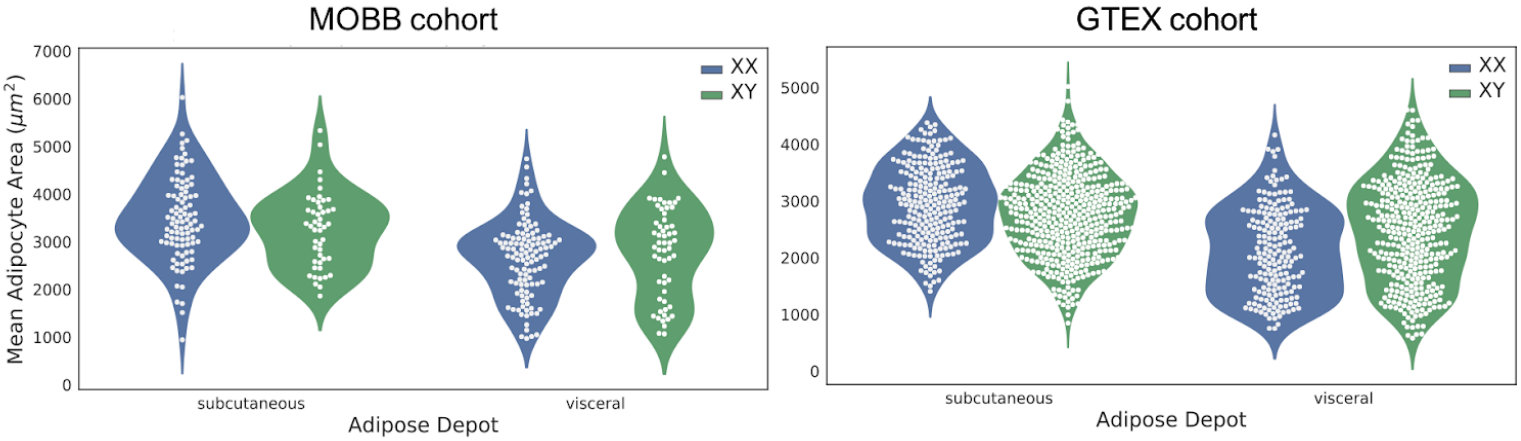


**Supplementary Figure 10 | Mean adipocyte area differences between adipose depots and sexes in GTEX and MOBB cohorts.** Females have significantly larger adipocytes in subcutaneous fat depots but smaller cells in visceral adipose tissue as compared to males.

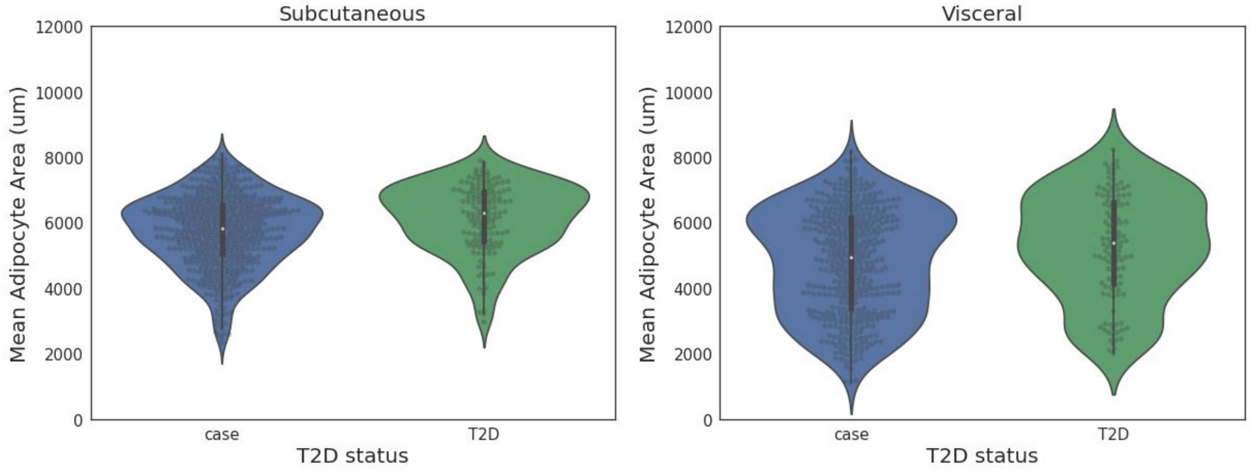


**Supplementary Figure 11 | Mean adipocyte area estimates for both subcutaneous and visceral adipose depots and their association with sample T2D status in the GTEx cohort.** Significant association with adipocyte cell size and T2D in both depots, which isn't significant after adjusting for BMI.

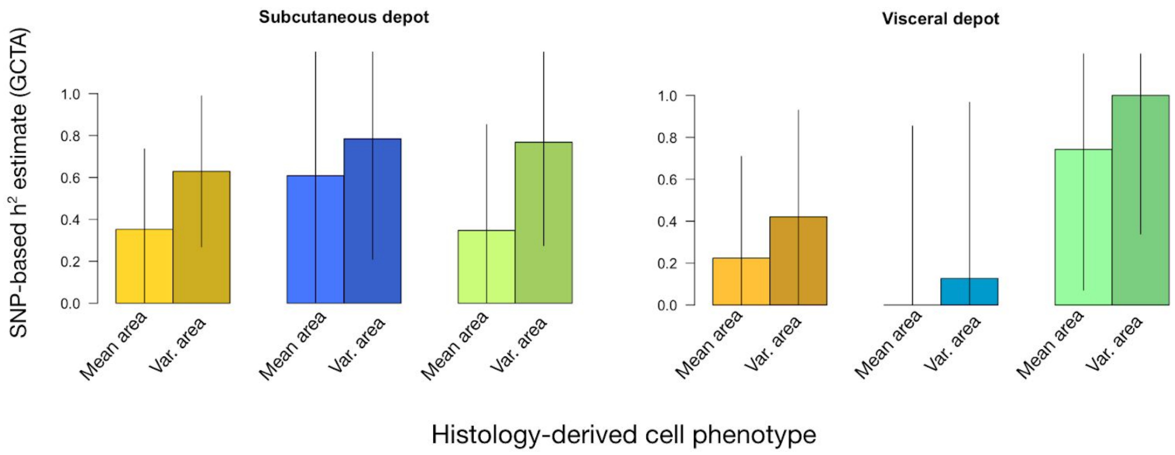


**Supplementary Figure 12 | Heritability estimates in histology-based phenotypes.** We estimated SNP-based heritability using Genome-wide Complex Trait Analysis (GCTA) in cell size mean (mean area) and cell size variance (var. area) derived from histology from both subcutaneous and visceral tissue. Heritability estimates range widely, and larger samples will be necessary to improve the precision of these estimates.

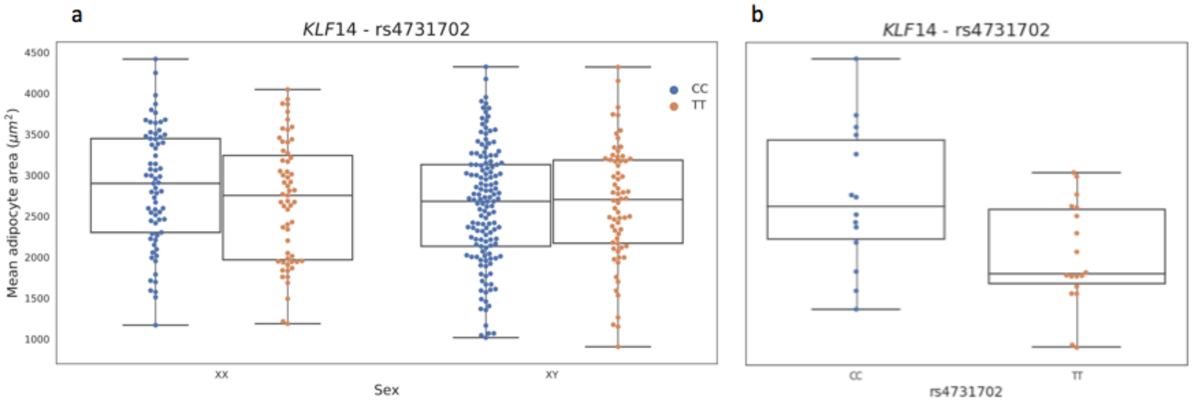


**Supplementary Figure 13 | Associations at the *KLF14* locus in GTEx samples.** (a) GTEx female rs4731702 risk allele (C) carriers have smaller adipocytes as compared to male carriers. (b) GTEx age and BMI restricted analysis show a significant association between rs4731702 and adipocyte size.

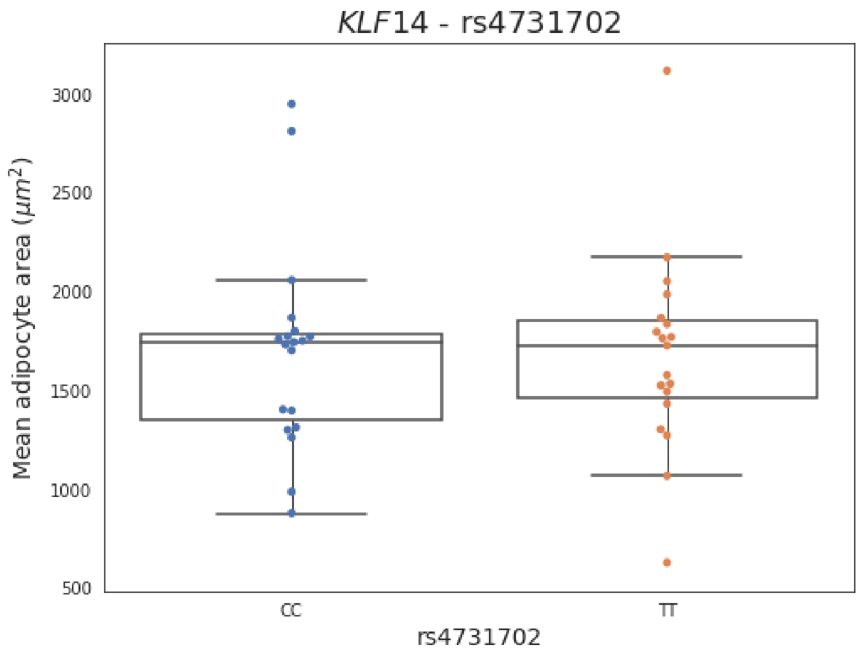


**Supplementary Figure 14 | Lack of association at the *KLF14* locus in** ENDOX **samples. (**a**)** female rs4731702 risk allele (C) carriers in ENDOX have no discernable difference in adipocyte size as compared to non-risk carriers (T) (CC = 19, TT = 19).

#### Supplementary Tables

| Depot | Sex | Phenotype | $h^2$ | SE |
|---|---|---|---|---|
| Subcutaneous | Combined | Mean cell area | 0.353 | 0.384 |
| | | Variance cell area | 0.629 | 0.360 |
| | Females | Mean cell area | 0.609 | 0.614 |
| | | Variance cell area | 0.785 | 0.575 |
| | Males | Mean cell area | 0.347 | 0.505 |
| | | Variance cell area | 0.768 | 0.494 |
| Visceral | Combined | Mean cell area | 0.224 | 0.485 |
| | | Variance cell area | 0.421 | 0.509 |
| | Females | Mean cell area | 0.000 | 0.854 |
| | | Variance cell area | 0.127 | 0.840 |
| | Males | Mean cell area | 0.742 | 0.671 |
| | | Variance cell area | 1.000 | 0.661 |

**Supplementary Table 1 | Heritability estimates for mean cell area and variance of cell area in both subcutaneous and visceral adipose tissue.** We used GCTA to estimate heritability in our largest cohort (GTEx). Results indicate that these cell morphology phenotypes are likely heritable. However, the errors are large, indicating that larger samples will be necessary to more accurately estimate heritability in these phenotypes.

**Supplementary Table 2 | Look-up of SNPs reported to associate with adipocyte morphology phenotypes.** We looked up rs4731702 and rs1421085, two SNPs previously reported to be associated with adipocyte morphology phenotypes, in our own genome-wide association results. We find no evidence for association at either SNP ($p > 0.05$ for all phenotypes and all sample groups). Results are provided as a text file in the accompanying GitHub repository:

rs4731702.cohort-level.gwas.results.txt
rs1421085.cohort-level.gwas.results.txt

**Supplementary Table 3 | Effect of obesity genetic risk scores on adipocyte area in μm$^2$ in different fat depots.** Estimates derived from a linear regression model with estimates of adipocyte area in μm$^2$ per 1-unit higher genetic risk score, corresponding to a predicted 1 standard deviation higher obesity trait and adjusting for 10 principal components in all cohorts. In GTEx, fatDIVA and MOBB, adjustments were also made for age and sex, and with and without adjusting for BMI. In the all-female ENDOX cohort, adjustments were made for 10 principal components and with and without adjusting for chip type as the other covariates were unavailable. In the main meta-analysis model (assuming a fixed-effect model), ENDOX was included adjusted for chip type. Two sensitivity meta-analyses were performed; including ENDOX unadjusted for chip type and excluding the ENDOX cohort.

mu.area.linear.regression.results.190305.xlsx

**Supplementary Table 4 | Effect of obesity genetic risk scores on standardized adipocyte area in different fat depots.** Estimates derived from a linear regression model of adipocyte area (standardized through rank inverse normal transformation of the residuals) per 1-unit higher genetic risk score, corresponding to a predicted 1 standard deviation higher obesity trait. Rank inverse normal transformations of the residuals were performed adjusting for 10 principal components in all cohorts. In GTEx, fatDIVA and MOBB, adjustments in the rank inverse normal transformation of the residuals were also made for age and sex, and with and without adjusting for body mass index. In the all-female ENDOX cohort, adjustments in the rank inverse normal transformation of the residuals were made for 10 principal components and with and without adjusting for chip type as the other covariates were unavailable. In the main meta-analysis model (assuming a fixed-effect model), ENDOX was included adjusted for chip type. Two sensitivity meta-analyses were performed; including ENDOX unadjusted for chip type and excluding the ENDOX cohort.

mu.area.linear.regression.results.190305.xlsx
